# Vesicular Trafficking Permits Evasion of cGAS/STING Surveillance During Initial Human Papillomavirus Infection

**DOI:** 10.1101/2020.03.29.014118

**Authors:** Brittany L. Uhlorn, Robert Jackson, Shauna M. Bratton, Shuaizhi Li, Koenraad Van Doorslaer, Samuel K. Campos

## Abstract

Oncogenic human papillomaviruses (HPVs) replicate in differentiating epithelium, causing 5% of cancers worldwide. Like most other DNA viruses, HPV infection initiates after trafficking viral genome (vDNA) to host cell nuclei. Cells possess innate surveillance pathways to detect microbial components or physiological stresses often associated with microbial infections. One of these pathways, cGAS/STING, induces IRF3-dependent antiviral interferon (IFN) responses upon detection of cytosolic DNA. Virion-associated vDNA can activate cGAS/STING during initial viral entry and uncoating/trafficking, and thus cGAS/STING is an obstacle to many DNA viruses. HPV has a unique vesicular trafficking pathway compared to many other DNA viruses. As the capsid uncoats within acidic endosomal compartments, minor capsid protein L2 protrudes across vesicular membranes to facilitate transport of vDNA to the Golgi. L2/vDNA resides within the Golgi lumen until G2/M, whereupon vesicular L2/vDNA traffics along spindle microtubules, tethering to chromosomes to access daughter cell nuclei. L2/vDNA-containing vesicles likely remain intact until G1, following nuclear envelope reformation. We hypothesize that this unique vesicular trafficking protects HPV from cGAS/STING surveillance. Here, we investigate cGAS/STING responses to HPV infection. DNA transfection resulted in acute cGAS/STING activation and downstream IFN responses. In contrast, HPV infection elicited minimal cGAS/STING and IFN responses. To determine the role of vesicular trafficking in cGAS/STING evasion, we forced premature viral penetration of vesicular membranes with membrane-perturbing cationic lipids. Such treatment renders a non-infectious trafficking-defective mutant HPV infectious, yet susceptible to cGAS/STING detection. Overall, HPV evades cGAS/STING by its unique subcellular trafficking, a property that may contribute to establishment of infection.

**Importance:** Persistent infection is the main risk factor for all HPV-associated cancers. However, cellular innate immune pathways exist to detect and limit viral infections. The cGAS/STING pathway senses cytosolic DNA to initiate antiviral IFN responses. Such responses would likely be detrimental towards the establishment of persistent HPV infections. We therefore hypothesize that HPV evades cGAS/STING detection via its unique vesicular trafficking mechanism. Here, we show that indeed HPV is a stealthy virus, capable of infecting keratinocytes with minimal activation of the cGAS/STING pathway. Such evasion is dependent on HPV’s vesicular trafficking, as perturbation of vesicular integrity during infection results in sensing of virions.

## Introduction

Human papillomaviruses (HPVs) are circular, double-stranded DNA viruses that infect and replicate in differentiating epithelium. HPV is the most commonly transmitted sexual infection, and oncogenic HPVs cause nearly 5% of all cancers worldwide and essentially all cervical cancers in women (1). The HPV lifecycle and viral gene expression is dependent on differentiating epithelium (2-4). To infect, HPV must traffic its circular dsDNA viral genome (vDNA) to the nucleus of basal keratinocytes, the only actively dividing cells within differentiating epithelium. Infected basal keratinocytes maintain vDNA episomes and serve as a reservoir for infected suprabasal cells that support the productive amplification of vDNA and assembly of progeny virions in upper epithelial layers (5, 6). Thus, persistently infected basal keratinocytes support sustained production of progeny virions to ensure efficient transmission within hosts. Notably, persistence of high risk HPV types is considered the risk factor for HPV-associated malignancies (7). Unraveling the mechanisms that underlie HPV persistence is therefore important to understand both HPV lifecycle and cancer.

Structurally, HPV is a simple virus, yet it has evolved a unique mitosis-dependent mode of subcellular trafficking and delivery of vDNA to the host cell nucleus (8-10). HPV virions bind to the extracellular matrix and cell surface heparan sulfate proteoglycans of basal keratinocytes, where upon proteolysis of the L1 and L2 capsid (11-13) and capsid conformational changes, the virion associates with entry receptor complexes and is endocytosed by the host cell (14-20). As the internalized virion travels through endosomal compartments, most of the L1 capsid is disassembled and degraded en route to the lysosome (9, 21, 22). In complex with the vDNA, minor capsid protein L2 facilitates retrograde trafficking of the vesicular vDNA away from the degradative endo/lysosomal compartment to the lumen of the Golgi, where it resides during interphase (23-27). L2 is an inducible transmembrane protein that can extend into the cytosol while remaining in complex with luminal vDNA by utilizing a transmembrane-like domain to protrude across local vesicular membranes to physically recruit retromer and other cytosolic sorting factors (8, 28-32).

Upon entry into mitosis, vesicular L2/vDNA traffics away from the fragmenting Golgi and accumulates on metaphase chromosomes via a chromatin-binding domain within the central portion of L2 (33). This L2-dependent chromosomal tethering of vDNA-containing vesicles ensures that the vDNA will be partitioned to the daughter cell nuclei after nuclear envelope reassembly and mitotic exit into G1. Our prior work using an L2-BirA fusion virus to report on L2/vDNA compartmentalization within vesicular membranes suggested that L2/vDNA may penetrate the Golgi-derived limiting membranes upon chromosome binding during prometa/metaphase to fully translocate into the nuclear/cytosolic milieu of open mitosis (27, 33). However, work from the Sapp and Schiller laboratories suggests that L2/vDNA remains vesicular through the completion of mitosis (34, 35). Recent work also shows that some L1 pentamers retrograde traffic along with the L2/vDNA complex towards the nucleus, but their role is unclear (36, 37). Additional data suggests that some virions may even remain somewhat intact as partially disassembled capsids during Golgi and post-Golgi nuclear trafficking (34). Within the daughter cell nuclei L2/vDNA somehow escapes the confines of these post-Golgi vesicles and recruits the nuclear ND10/PML body components necessary for efficient early viral gene expression (34, 35, 37, 38).

Several other non-enveloped viruses, such as adenoviruses, nodaviruses, parvoviruses, picornaviruses, and reoviruses, have evolved more direct means of penetrating limiting membrane barriers for vDNA/vRNA delivery and subsequent infection of host cells (39-41). The evolutionary rationale behind the unique mitosis-dependent vesicular trafficking mechanism of HPV is unknown. We and others speculate the trafficking mechanism may impart immunoevasive properties via vDNA shielding behind protective limiting membranes en route to the nucleus (8, 34, 42, 43).

The innate immune system is adept at recognizing microbial nucleic acids as pathogen-associated molecular patterns (PAMPs) through a number of pattern recognition receptors (PRRs) (44). Many PRRs also function to sense nucleic acids of cellular origin as danger-associated molecular patterns (DAMPs). These DAMPs typically consist of mislocalized or modified nucleic acids within the context of physiological stresses or cellular damage often associated with microbial infections (45, 46). In general, the activation of PRRs by nucleic acids leads to NFκB-dependent inflammatory cytokine responses and/or IRF3/7-dependent type-I interferon (IFN) antiviral responses.

The cGAS/STING pathway has emerged as a central innate immune sensing pathway for cytosolic DNA and downstream IFN responses (47-52). Briefly, cytosolic DNA is recognized by the enzyme cGAS, triggering production of the cyclic dinucleotide 2’,3’-cGAMP (53, 54). STING, a transmembrane endoplasmic reticulum (ER) protein (55), is activated upon binding to 2’,3’-cGAMP (56). Once activated by 2’,3’-cGAMP, dimeric STING traffics to a perinuclear Golgi-like compartment (57, 58), where it oligomerizes to recruit and activate TBK1 (59-61) to phosphorylate the transcription factor IRF3, stimulating an IFN response (55).

Initial entry, trafficking, and uncoating of many DNA viruses like herpesviruses (HSV-1, KSHV, HCMV), poxviruses (VACV, MVA), asfarviruses (ASFV), and adenoviruses (HAdV serotypes 2, 5, 7, and 35) have been shown to activate cGAS/STING (62-69). Cellular sensing of initial HPV infection has not been formally investigated, but several studies have described roles for the early HPV oncogenes in antagonizing the cGAS/STING/IRF3 axis upon establishment of infection (70-72). Active antagonism of cGAS/STING by the early HPV proteins suggests that cGAS/STING responses would oppose persistent HPV infection. Indeed, IFN responses are detrimental to persistent HPV infections, reducing cellular proliferation and causing apoptosis, episome loss, mutation, and/or integration (73-77). Likewise, it is well known that some HPV early genes – E5, E6, and E7 – counteract these detrimental antiviral IFN and IFN-stimulated gene (ISG) responses through a variety of mechanisms (78-81).

Given the ongoing interplay between HPV and antiviral responses, we postulate that evasion of cGAS/STING responses during initial infection would be beneficial to the viral lifecycle. We hypothesize that HPV’s unique mitosis-dependent vesicular trafficking serves to enable such evasion. Here, we directly compare human keratinocyte cGAS/STING responses to HPV16 virion infection versus responses to cationic lipid transfection of dsDNA plasmid. We show a striking lack of IRF3 phosphorylation upon HPV16 infection. RNA-seq confirmed the lack of downstream antiviral IFN and ISG transcriptional responses to HPV16 infection. Perturbance of intracellular vesicular membranes during infection results in antiviral responses to HPV16 infection, suggesting that HPV’s distinctive vesicular trafficking underlies its stealthy abilities.

## Results

### HPV16 Evades cGAS/STING Responses During Initial Infection

We initially investigated cellular cGAS/STING responses to HPV16 pseudovirus in HaCaT cells, spontaneously immortalized keratinocytes (82) that have a functional DNA-responsive cGAS/STING pathway (83-85). HaCaT cells were either transfected with 250 ng of endotoxin free dsDNA plasmid (pGL3) for 90 minutes or infected with the HPV16 virion equivalent to 250 ng DNA (approximately equivalent to 950 ng L1 capsid, encapsidating pGL3). Cellular responses to DNA transfection were followed over the next 12 hours by western blotting. Since HPV16 infection is slow and asynchronous (14) with different DNA delivery kinetics compared to transfection, HaCaT cells were infected with HPV16 pseudovirus and cGAS/STING activity was assessed over an extended 24 hour time course. In these experiments, virus was present until the sample was collected for analysis. An acute and robust IRF3 phosphorylation (pIRF3) was observed in response to DNA transfection (Figure 1A, left panel). In contrast, robust phosphorylation of IRF3 was not evident at any time point post-infection with HPV16 pseudovirus, indicating that the cGAS/STING pathway was not able to sense and respond to HPV16 vDNA (Figure 1A, B).

**Figure 1.**
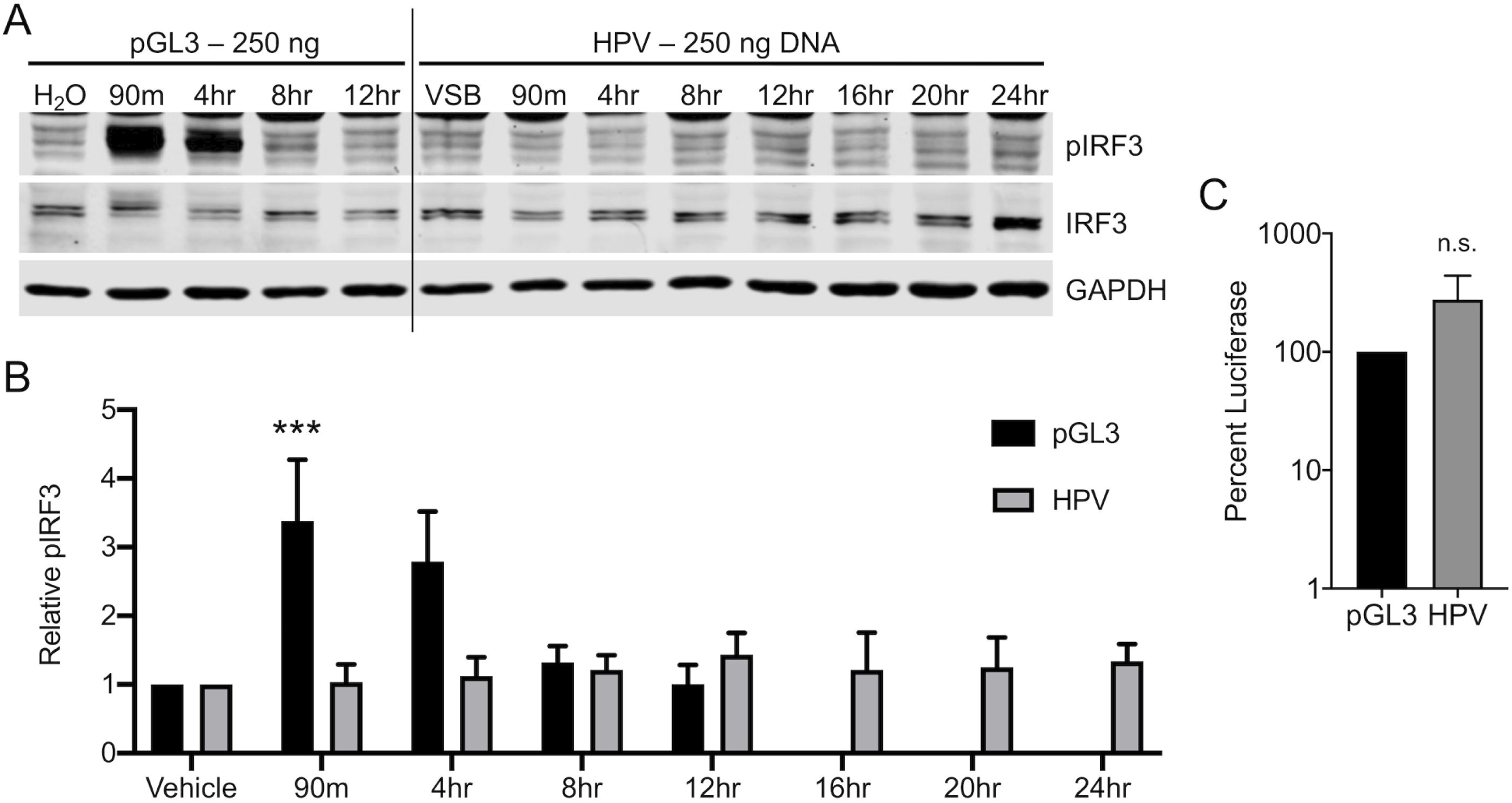
HPV16 Evades cGAS/STING Responses in HaCaTs During Initial Infection. cGAS/STING responses to pGL3 and wildtype PSV in HaCaTs. **(A, B)** Cells were either transfected for 90 min with 250 ng pGL3 or infected for the duration of the experiment with 950 ng L1 (equivalent to 250 ng DNA) of HPV16 virions. Cells were harvested at various times post-treatment and cGAS/STING responses assessed via western blotting. **(A)** IRF3 was phosphorylated, most prominently at 90 min and 4 hr post pGL3 transfection, while IRF3 was not phosphorylated at any time post HPV16 infection, representative blot of *n* = 4 biological replicates. **(B)** Densitometric quantification of western blots, *n* = 4 biological replicates. Statistics calculated by two-way ANOVA (*P*_*interaction*_ = 0.0138) followed by Sidak’s multiple comparison test (\*\*\**P* < 0.005). **(C)** Cells were either transfected for 90 min with 250 ng pGL3 or infected for the duration of the experiment with an equivalent amount of HPV16 virions. Luciferase activity was measured 24 hr post-treatment, normalized to GAPDH, and corrected for pGL3 encapsidation as described in *Materials and Methods*. Differences were not significant (n.s.) by unpaired *t*-test, *n* = 3 biological replicates.

These HPV16 virions package pGL3, which contains a firefly luciferase expression cassette driven by an SV40 promoter. To gauge the relative pGL3 delivery between plasmid transfection and HPV16 infection, luciferase assays were performed 24 hours post-treatment. Since papillomaviruses lack specific genome packaging signals (86), HPV16 pseudoviruses generated in the 293TT system will package both pGL3 reporter plasmid and cellular dsDNA ≤ 8kb in size. The majority of virions will contain 293TT-derived DNA, with a smaller fraction containing pGL3 (87). For simplicity, the encapsidated DNA will be referred to as “vDNA” herein, even though it is not the authentic viral genome. To account for partial packaging of pGL3, the luciferase numbers for the HPV16 infections were corrected by factoring in the measured virion:pGL3 ratio of 11.16 (See *Materials and Methods*). Despite a complete lack of pIRF3 responses, HPV16 infection resulted in equivalent luciferase activity compared to pGL3 transfected cells (Figure 1C), arguing that differences in cellular DNA delivery do not explain the apparent evasion of cGAS/STING surveillance by HPV16.

While immortalized HaCaT cells represent a good model for the basal keratinocytes that HPV has tropism for *in vivo*, we sought to investigate cGAS/STING responses in cultured primary human foreskin keratinocytes (HFKs), a better model for the cells HPV naturally infects. Primary HFKs were transfected with 500 ng pGL3 or infected with the viral DNA equivalent. pGL3 transfection resulted in robust activation of the cGAS/STING pathway, as seen by the detection of phosphorylated STING (pSTING) and pIRF3 (Figure 2). As before, HPV16 infection did not activate the pathway as detected by phosphorylation of STING or IRF3 during infection (Figure 2A). We observed a delay in activation between HaCaTs (peak near 90 minutes) and HFKs (4 hours). This may be due to differences in transfection efficiency between primary HFKs and HaCaTs, or the kinetics of cellular activation may be slower in primary cells. To ensure comparable delivery of the 500 ng dsDNA via HPV16 virions compared to plasmid transfection we performed a luciferase assay 24 hours post-transfection/infection. As with HaCaT cells, HPV16 infection again yielded robust luciferase activity as compared to plasmid transfection of HFKs (Figure 2B). These data further support a stealthy mode of HPV infection whereby cGAS/STING responses are avoided.

**Figure 2.**
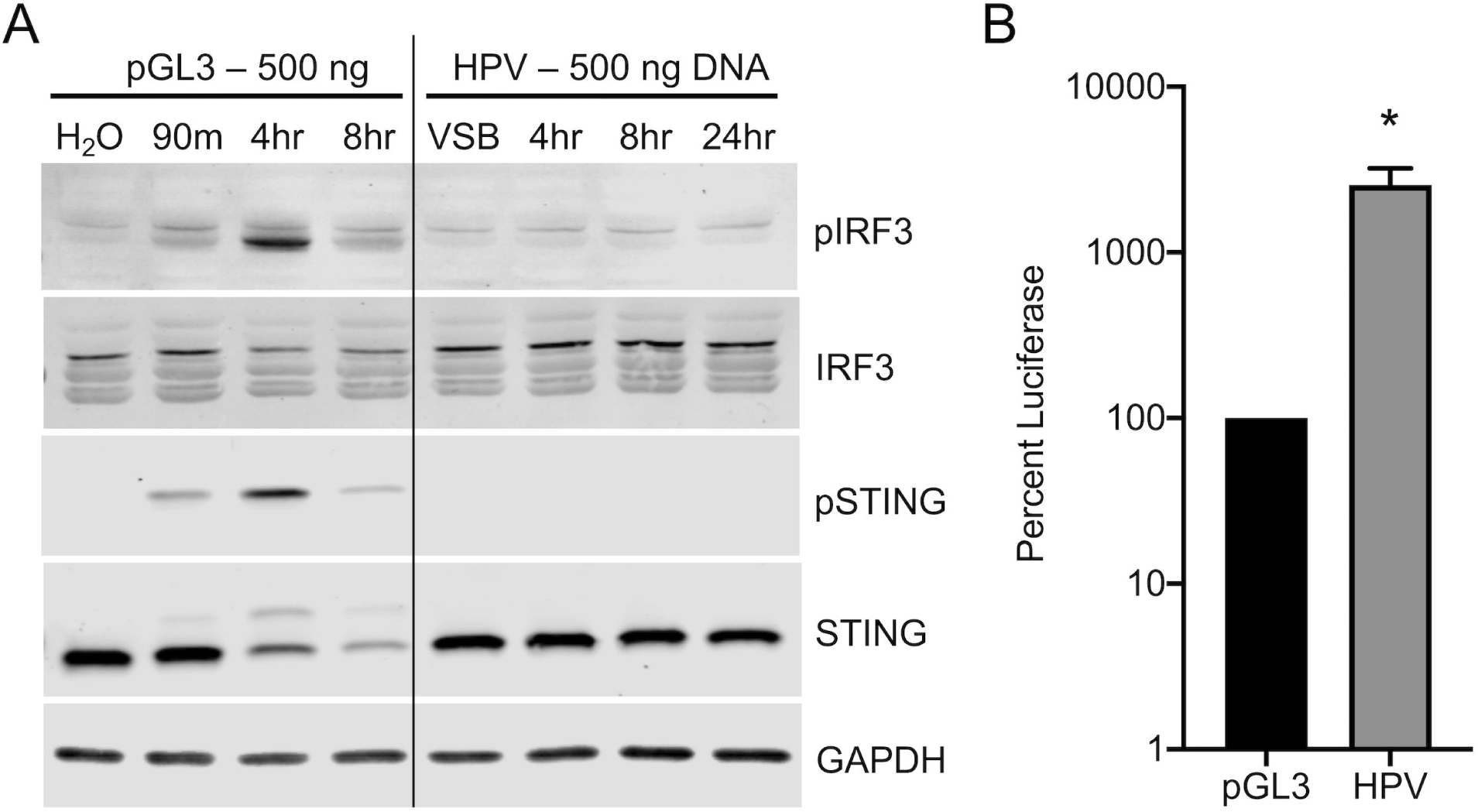
HPV16 Evades cGAS/STING Responses in HFKs During Initial Infection. cGAS/STING responses to pGL3 and wildtype HPV16 in primary HFKs. **(A)** HFKs were either transfected for 90 min with 500 ng pGL3 or infected for the duration of the experiment with 1900 ng L1 (equivalent to 500 ng DNA) of HPV16 virions. Cells were harvested at various times post-treatment and cGAS/STING responses assessed via western blotting. **(A)** IRF3 was phosphorylated 4 hr post pGL3 transfection, while IRF3 was not phosphorylated at any time post HPV16 infection with HPV PsV. representative blot of *n* = 3 biological replicates. **(B)** Cells were either transfected for 90 min with 250 ng pGL3 or infected for the duration of the experiment with an equivalent amount of HPV16 virions. Luciferase activity was measured 24 hr post-treatment, normalized to GAPDH, and corrected for pGL3 encapsidation as described in *Materials and Methods*. \**P* < 0.05 by unpaired *t*-test, *n* = 3 biological replicates.

### Transcriptional Responses to DNA Transfection and HPV16 Infection of Primary HFKs

We performed RNA-seq analysis to profile cellular transcriptional responses to dsDNA delivered via cationic liposomes or HPV16 pseudovirions (Figure 3, Data Set S1, S2). As above, HFKs were either transfected with 500 ng pGL3 or infected with the 500 ng DNA equivalent of HPV16 virions. Total RNA was extracted at the indicated timepoints. At 4 hours post-transfection, 142 genes (95 up and 47 down) were differentially expressed (Figure 3A, Data Set S3) in HFKs relative to mock water transfection (*n* = 2, *P*-adj < 0.05). The up-regulated genes (Figure 3B), representing expected transcriptional responses to cytosolic DNA, were functionally enriched (see Data Set S4 for detailed results) for 82 Gene Ontology (GO) Biological Processes, including “defense response to virus” (GO:0051607, *P*-adj = 2.26 × 10^−15^) as the top hit among innate immune-related terms. Of the 10 enriched Reactome Pathways, the top hit was “Interferon alpha/beta signaling” (REAC:R-HSA-909733, *P*-adj = 1.16 × 10^−7^). In line with our western blot data (Figures 1 and 2), IRF3 was the top transcription factor signature related to these up-regulated genes (*P*-adj = 8.61 × 10^−10^). The top five most significantly up-regulated genes at 4 hours post-transfection included *IFIT2, IFIT3, ATF3, NOCT*, and *IRF1*; the genes with the highest fold change, ranging from ∼70 to 400 times above baseline, were *IFNL1, IFNL2, IFIT2, NEURL3*, and *ZBTB32*.

**Figure 3.**
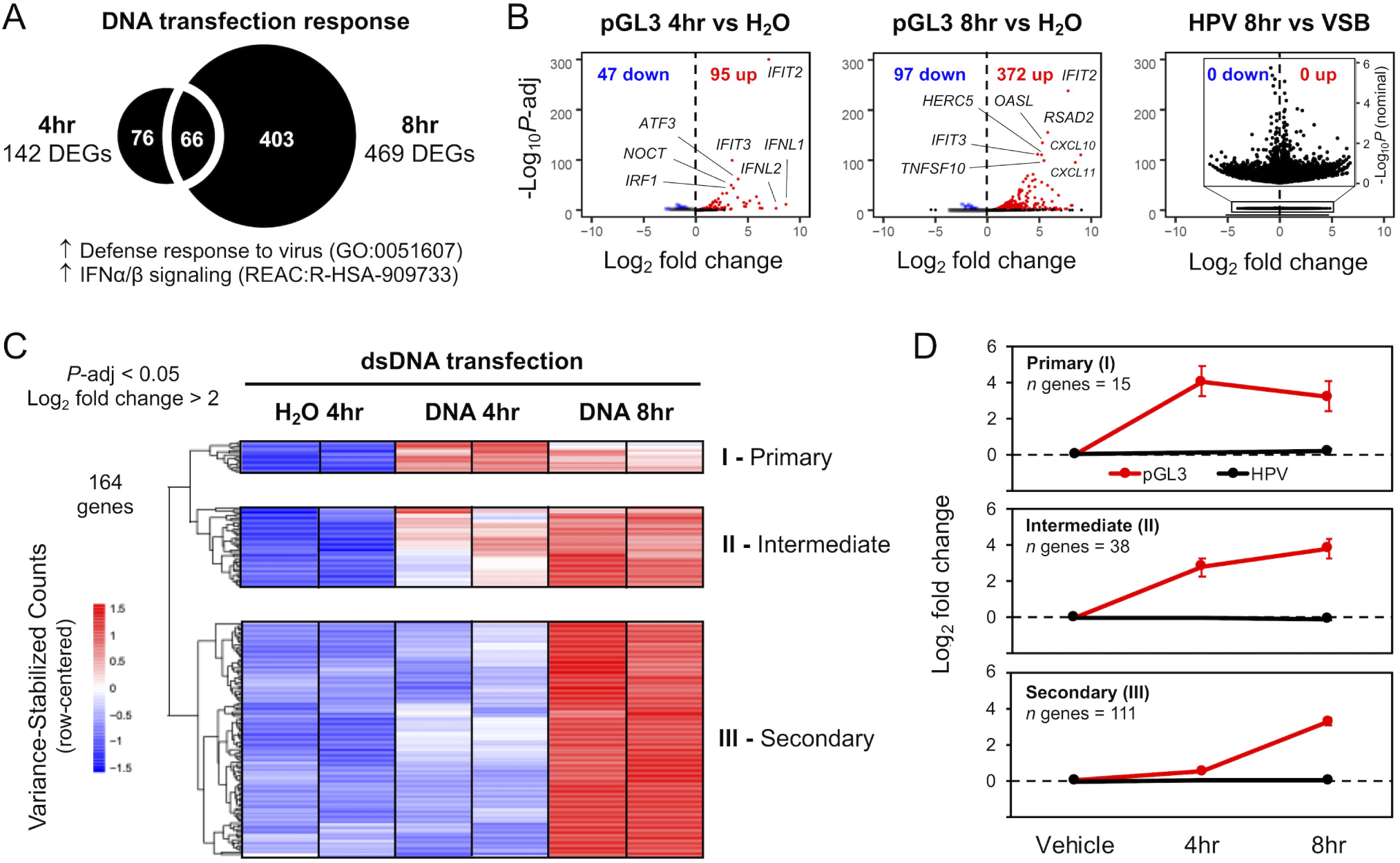
HPV16 Virion Infection Does Not Activate Cellular Responses to dsDNA. RNA-seq was used to transcriptionally profile cellular responses to pGL3 DNA introduced via liposome transfection or HPV virion infection. **(A)** Relative to H_2_O mock-transfection, there were 142 significantly (*P*-adj < 0.05) differentially expressed genes (DEGs) at 4hr and 469 DEGs at 8hr post-transfection, with up-regulated genes functionally enriched for biological processes and pathways related to defense responses to virus and IFN signaling. **(B)** Volcano plots for 4 and 8hr pGL3 transfections (*n* = 2) show significantly DEG genes (*P*-adj < 0.05), with up-regulated genes (red) related to strong innate immune signaling responses. No response was activated 8hr following HPV virion infection (*n* = 3) relative to viral storage buffer (VSB); inset volcano plot is re-scaled, using nominal (unadjusted) *P*-values, to show distribution of genes that do not reach significance. **(C)** Heatmap of variance-stabilized counts, row-centered, for the top up-regulated cellular response genes (*P*-adj < 0.05, log_2_ fold change > 2, for 4 and 8hr pGL3-transfection: 164 genes). Genes clustered into three main groups, corresponding to temporal responsiveness post-transfection: cluster I (immediate/primary response at 4hr), cluster II (intermediate response at 4 and increasing at 8 hr), and cluster III (secondary response at 8 hr). **(D)** Metagene log_2_ fold change values were computed by aggregating response data of all genes within each cluster. Error bars represent 95% confidence intervals for each gene cluster.

At 8 hours post-transfection, 469 genes (372 up and 97 down) were differentially expressed (Figure 3A, Data Set S3) in HFKs (*n* = 2, *P*-adj < 0.05). These up-regulated genes were also functionally enriched for IFN signaling, with the top five most significantly up-regulated genes (Figure 3B) at 8 hours post-transfection including *IFIT2, RSAD2, OASL, IFIT3*, and *HERC5*. The genes with the highest fold change at 8 hours, ranging from ∼200 to 500 times above baseline, were *CXCL10, CXCL11, IFNL1, GBP4*, and *IFIT2*. Investigation of the heatmap (Figure 3C) identifies 3 main functional groups of genes (Figure 3D). Genes that are upregulated acutely, at 4 hours (I; ‘Immediate’/’Primary’), and either go back down (‘Immediate’) or remain up at 8 hours (‘Primary’), genes that are upregulated by 4 hours and continue to increase throughout the experiment (II; ‘Intermediate’), and a group of genes that are upregulated by 8 hours but were not significantly affected at the 4 hour timepoint (III; ‘Secondary’). Complete gene sets for each cluster are provided in supplementary Data Set S5.

While 500 ng of pGL3 delivered via cationic liposomes triggered a robust IRF3-based transcriptional response in HFKs, with the strongest overall response at 8 hours post-transfection, the equivalent amount of DNA delivered via HPV16 pseudovirus infection yielded no response (Figure 3B, right panel). At 8 hours post-infection, there were no genes differentially expressed in HFKs relative to mock infection with viral storage buffer (*n* = 3). Importantly, preliminary analysis of later time-points did not show activation of IFN or ISG related transcription (Figure S1). Overall, HPV infection is unable to induce a classic innate immune response signature in primary HFKs (Figure 3B, D).

Figure 4 highlights the transcriptional response of different IFN and ISGs that were previously demonstrated to interact with the HPV lifecycle (see *Discussion*). These responses fall into the same functional classes identified in Figure 3: 1) an acute response peaking at 4 hours (e.g., *IFNB1, IFNL1, IFNL2, ATF3*, and *IRF1*), 2) an intermediate response with an apparent peak at 8h (e.g., IFIT1, HERC5, IRF7, ISG15, and *CDKN2C*), and 3) genes belonging to class III, displaying a ‘secondary’ response (e.g., *TNFSF10, RSAD2, IFI16, CXCL10*, and *CXCL11*). Similar to Figure 3, there is no evidence (Figure 4) that HPV16 infection upregulates the transcription of these genes.

**Figure 4.**
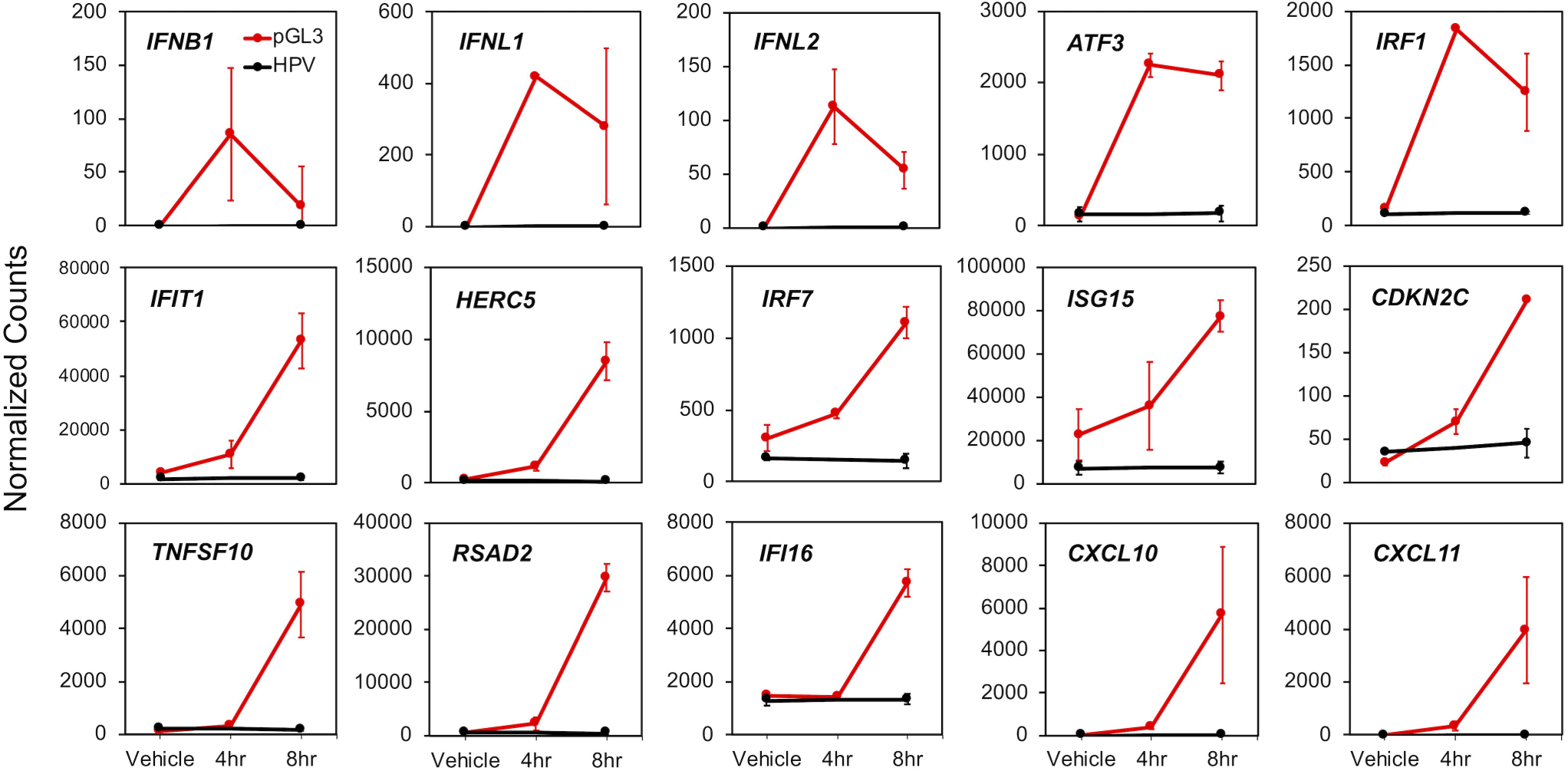
HPV-related DNA-responsive cellular genes are not induced by virion infection. Normalized RNA-seq count data were plotted for a subset of 15 DNA-responsive cellular genes selected based on prior literature indicating a relationship with HPV. Error bars represent 95% confidence intervals (*n* = 2 for pGL3 transfection, red, and *n* = 3 for HPV virion infection, black).

### Bypassing HPV’s Natural Trafficking Pathway Activates cGAS/STING

cGAS/STING is capable of sensing the incoming vDNA from a number of viruses including adenoviruses, poxviruses, herpesviruses, and the reverse-transcribed cDNA products of lentiviruses like HIV (62, 63, 88, 89). These viruses either breach (non-enveloped viruses) or fuse with (enveloped viruses) intracellular limiting membranes to provide vDNA-capsid complexes access to the cytosol and eventually reach the nucleus. In contrast, HPV uses minor capsid protein L2 to transport vDNA within vesicular membranes en route to the nucleus during mitosis-dependent subcellular trafficking (8-10). We hypothesize this unique vesicular trafficking enables evasion of cellular cGAS/STING surveillance.

To test this hypothesis, we stimulated premature membrane penetration of HPV vDNA to determine if cGAS/STING would then be capable of detecting HPV infection. As no HPV capsid protein mutations are known to cause leakage or transfer of vDNA across vesicular membranes, we instead used cationic lipids which are known to perturb intracellular limiting membranes (90-92). Indeed, our prior work using a sensitive enzyme-based L2 membrane penetration assay showed that even low amounts of cationic lipids during HPV infection are capable of disrupting vesicular membranes that normally limit HPV from cytosolic exposure (27).

HPV16 virions encapsidating pGL3 were mixed to the cationic lipid Lipofectamine 2000, as described in *Materials and Methods*. Transmission electron microscopy of this mixture revealed that intact HPV16 virions were in the close vicinity of submicron cationic lipid complexes (Figure 5A). We next assessed the ability of wildtype HPV16 and the post-Golgi trafficking defective R302/5A mutant HPV16 to deliver vDNA to the nucleus in the presence of cationic lipids compared to control media. The R302/5A mutation in L2 renders particles non-infectious. Particles that contain R302/5A mutant L2 undergo normal early trafficking but fail to penetrate Golgi-derived vesicular compartments due to defective binding between the mutant L2 and host mitotic chromosomes (27, 33, 93). R302/5A vDNA therefore remains cloaked within vesicular membranes, failing to reach the nucleus. Addition of cationic lipids caused a modest ∼60% decrease in wildtype HPV16 infectivity, as measured by luciferase assay for the delivery of virion-packaged pGL3 to the nucleus (Figure 5B). In stark contrast, cationic lipids rescued the infectivity of R302/5A mutant virus to levels comparable to that of wildtype HPV16 in the presence of cationic lipids (Figure 5B). The natural subcellular retrograde trafficking of vDNA is dependent on endosomal acidification, furin cleavage of L2, and γ-secretase activity. Small molecule compounds that perturb these processes are potent inhibitors of HPV16 infection (8). The cationic lipid-dependent infectivity of R302/5A was insensitive to endosomal acidification inhibitor BafA, furin inhibitor dRVKR, or γ-secretase inhibitor XXI (Figure 5C). Collectively these data indicate that cationic lipids mediate vesicular escape of HPV16 to rescue infectivity of R302/5A via an alternative route, independent of the classical HPV16 vesicular trafficking pathway.

**Figure 5.**
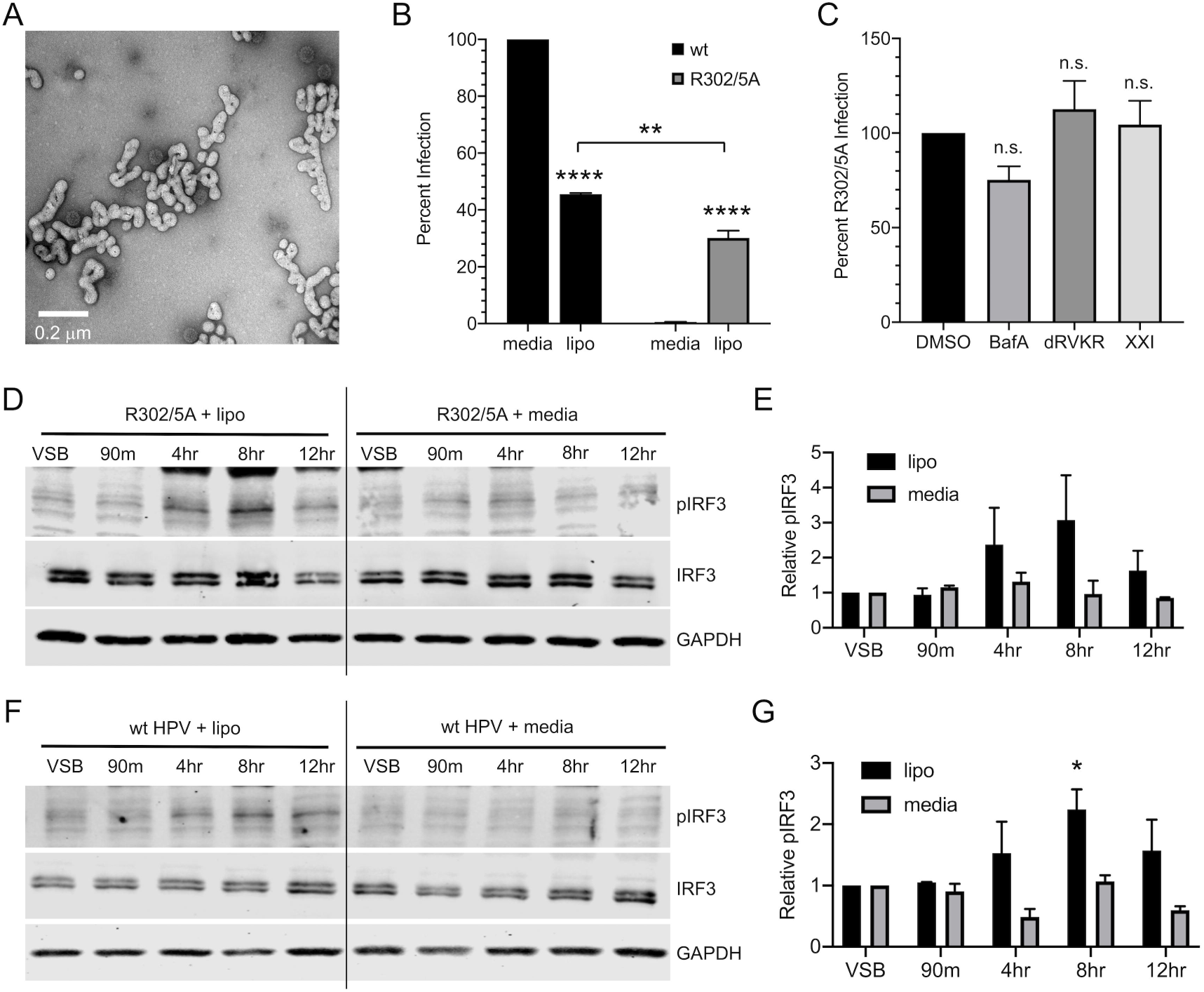
Bypassing HPV’s Natural Trafficking Pathway Activates cGAS/STING. Addition of cationic lipids during infection with translocation-deficient R302/5A mutant HPV16 restores infectivity and allows for cGAS/STING sensing. **(A)** Electron micrograph of HPV16 complexed with the cationic lipid Lipofectamine 2000. **(B)** HaCaTs were infected with 2e8 vge/well of wildtype or R302/5A virion +/- cationic lipid (lipo) for 4 hr and infection was measured by luciferase assay 24h post-infection. While naturally non-infectious, addition of cationic lipids restored infectivity of the R302/5A mutant to levels nearly comparable to those of wildtype HPV16. Statistics calculated by one-way ANOVA (*P* < 0.001) followed by Tukey’s multiple comparisons test (\*\**P* < 0.01, \*\*\*\**P* < 0.001), *n* = 3 biological replicates **(C)** R302/5A infection in the presence of cationic lipids is insensitive to biochemical inhibitors of endosomal acidification (BafA), furin (dRVKR), and γ-secretase (XXI). Differences were not significant by one-way ANOVA followed by Dunnett’s multiple comparisons test, *n* = 3 biological replicates. **(D-G)** Cells were infected for 4 hr with R302/5A **(D, E)** or wildtype HPV16 **(F, G)** virion equivalent of 250 ng DNA +/- cationic lipids (lipo) as described in *Materials and Methods*. Addition of cationic lipids during virus infection resulted in IRF3 phosphorylation at 4 hr and 8 hr, while virus infection in media alone did not induce IRF3 phosphorylation, representative blots shown of *n* = 2 biological replicates in both **(D, F). (E, G)** Densitometric quantification of western blots. **(E)** Statistics were calculated by two-way ANOVA followed by Dunnett’s multiple comparisons test. Due to technical aspects of background variability and few replicates, differences did not reach statistical significance (*P*_*interaction*_ = 0.331, *P* = 0.125 at 8 hr time point), *n* = 2 biological replicates. **(F)** Statistics were calculated by two-way ANOVA (*P*_*interaction*_ = 0.1359), followed by Dunnett’s multiple comparisons test (\**P* < 0.05), *n* = 2 biological replicates.

Since the use of cationic lipids to bypass the natural subcellular trafficking pathway of HPV rescued infectivity of R302/5A mutant virions, we examined whether premature disruption of vesicular membranes by cationic lipids would allow for activation of the cGAS/STING pathway by HPV16. HaCaT cells were infected with R302/5A virions +/- cationic lipids and cGAS/STING activity was assessed by pIRF3 blot as before. Similar to wildtype HPV16 infection (Figure 1), R302/5A mutant infection alone did not activate the cGAS/STING pathway. However, premature vesicle disruption allowed for robust sensing of DNA delivered by R302/5A infection as measured by IRF3 phosphorylation, most prominently at 4 and 8 hours post-infection (Figure 5D, E). Similar results were seen in experiments using wt HPV16 particles (Figure 5F, G). These results indicate that when cationic lipids allow HPV16 virions to breach vesicular compartments, the cGAS/STING pathway is activated, generating a pIRF3 response.

## Discussion

Cells are equipped with numerous PAMP- and DAMP-sensing PRRs, designed to detect the first signs of cellular stress and respond in an IRF3 and NFκB transcription factor-dependent manner to induce the expression of IFN, antimicrobial ISGs, and proinflammatory cytokine genes (94). Geared towards recognizing dsDNA in the cytoplasm of a cell, the recently characterized cGAS/STING pathway has proven to be a critical arm of innate immunity and an important cellular antiviral defense system that is targeted for evasion and/or antagonism by many different viruses (51, 52, 95).

The HPV early proteins have been shown to block cGAS/STING and downstream IRF3-dependent IFN responses, as well as NFκB-dependent cytokine responses through a variety of mechanisms (70, 71, 79-81, 96, 97). The early viral proteins that counteract the cellular antiviral responses are not packaged inside within incoming virions. Thus, evasion mechanisms to limit early detection by the innate immune system during early infection would likely benefit later stages of the viral lifecycle. Viral blunting of these cell intrinsic and extrinsic responses promote viral persistence by enabling maintenance of episomal vDNA and reducing inflammatory responses that cause activation of antigen-presenting cells and promote adaptive antiviral immune responses (98).

Here we investigate cGAS/STING responses to initial HPV16 infection. Although some prior work has reported minimal induction of IFN and downstream ISGs in response to HPV and canine PV infection (99, 100), no studies have directly addressed cGAS/STING responses to incoming HPV16 virions. Importantly, other than the viral L1 and L2 proteins that are part of the incoming viral capsid, the pGL3 reporter plasmid does not express any papillomavirus proteins. We find that while keratinocytes mount acute pIRF3 and downstream IFN/ISG responses to dsDNA plasmid transfection, dsDNA delivered through HPV16 infection proceeds undetected by cGAS/STING. To allow for direct comparison, we infected cells with the virion equivalent of 500 ng DNA, corresponding to a multiplicity of infection (MOI) of 400,000 virions/cell. These super-physiological conditions strengthen the hypothesis that a physiological infection is unlikely to be detected. The transcriptional response to dsDNA included type-I *IFNB1*, type-III *IFNL1* and *IFNL2*, and many IFN-dependent ISGs and chemokines (Figs. 3 and 4). Many of these ISGs, like *IFIT1/2/3, ISG15* and *CXCL10/11*, can be upregulated directly through IRF3-mediated transcription, in addition to secondary IFN-dependent signaling (pSTAT1/pSTAT2/IRF9) mechanisms in certain cellular contexts (101-103).

Not surprisingly, many of these proteins have been implicated in the viral lifecycle. *IFIT1* (p56/ISG56) has the potential to directly restrict persistent HPV infection via inhibition of E1-dependent episomal vDNA maintenance (77). *ISG15* encodes a ubiquitin-like molecule with broad antiviral activity. *HERC5* is an E3 ligase that catalyzes the addition of ISG15 to nascent viral proteins leading to the restriction of HPV replication (104). CXCL10, CXCL11, and related CXCL-family chemokines attract activated T cells via CXCR3 (105, 106). These chemokine responses may also be detrimental to HPV persistence, as evidenced by a recent mouse papillomavirus study where MmuPV1 was observed to specifically downregulate stress keratin-induced expression of CXCR3 ligands including CXCL10 (107). *IFI16* is a DNA-binding inflammasome component and can restrict HPV replication by epigenetically silencing viral gene expression reducing vDNA copy number (108, 109). Several of the genes upregulated by dsDNA transfection are counteracted by papillomavirus gene products. *IRF1* and *IRF7* gene products augment IFN/ISG responses. IRF1 activity is counteracted by the E7 protein from high risk HPVs (110, 111). Similarly, *RSAD2* (viperin) has broad antiviral activity and is downregulated by cutaneous HPV2 E7 (112).

Double-stranded DNA transfection also induced cell survival and proliferation modulators like *ATF3, CDKN2C*, and *TNFSF10. ATF3* interferes with E6’s ability to degrade p53 by preventing E6AP from binding to p53, thus interfering with HPV immortalization (113). Interestingly, *ATF3* expression is downregulated in cervical cancer (113). *CDKN2C* encodes p18INK4C, which restricts cellular proliferation through inhibition of CDK4 and CDK6 (114). Expression of *CDKN2C* is upregulated by oncogenic E6 proteins (115), suggesting that the virus must be able to replicate in the presence of this anti-proliferative signal. *TNFSF10* encodes TRAIL, a member of the TNF family of ligands that can initiate apoptosis (116). The viral E5 protein counteracts TRAIL signaling to block apoptosis and promote HPV infection (117, 118). If activated, the induction of cellular effectors would likely restrict initial HPV replication following infection. Thus, initial evasion of cGAS/STING and these downstream cellular IRF3/IFN/ISG responses would likely be beneficial to HPV infection. Collectively, our data suggest that the ability of the viral proteins to antagonize so many of these effectors is not to counteract sensing of the viral DNA upon initial infection. Rather, counteraction of these antiviral/antitumor responses is important for later steps in the viral lifecycle.

Interestingly, HPV16 infection is sensed when vesicular membranes are perturbed by the inclusion of cationic lipids during infection. These findings also indicate that the chromatinized encapsidated vDNA is insufficient to (completely) mask the viral genome from cGAS surveillance, as recent reports have shown that the nucleosomal components of mitotic chromatin prevents efficient cGAS activation by dsDNA (85, 119, 120). We conclude that HPV’s unique L2-dependent vesicular membrane trafficking effectively shields the vDNA PAMP from cytosolic cGAS/STING surveillance during nuclear transit, likely contributing towards viral persistence. A recent report describes a DNA-PK mediated DNA-sensing mechanism that appears to be specific to linear DNA (121). Since the HPV genome is circular, it is unlikely that this pathway would be involved in the sensing of HPV infection.

Our data suggest that papillomaviruses use subcellular trafficking to evade innate immune detection. Variation in subcellular trafficking routes has been shown to influence innate immune responses to other viruses. Different trafficking mechanisms during HAdV entry of several cell types have been shown to affect the stimulation of certain PRRs, influencing distinct immune responses that are dependent on particular virus- and cell-type combinations (122). For example, different HAdV serotypes naturally vary in their receptor usage as well as downstream early vs. late endosomal trafficking pathways. These inherent differences in HAdV serotypes have been shown to affect innate TLR-dependent sensing of the virus (123). For example, Dynamin-2 (DNM2) is a cellular GTPase involved in microtubule-dependent transport and scission of endosomes. Modulation of DNM2 affects the trafficking of HAdV, resulting in an altered cellular cytokine response to infection (124). Reovirus T1D and T3L strains elicit differing magnitudes of IRF3-dependent IFN responses in a manner dependent on viral uncoating during the late steps of viral entry and trafficking (125). In the case of HIV, the CD4 receptor and the cellular dynein adapter BICD2 play roles in viral trafficking, influencing innate sensing and downstream IFN/ISG responses in infected cells (126, 127). Likewise, host phospholipase D affects innate sensing of the influenza A virus by modulation of entry (128).

Thus, while differing modes of trafficking can affect cellular responses to many viruses, HPV appears to have evolved an extremely covert means of evading cellular IRF3-dependent responses by hiding incoming vDNA inside the vesicular trafficking network. Other viruses may have evolved similar trafficking-dependent immunoevasion strategies. Adeno-associated viruses (AAVs) are parvoviruses that have recently been shown to retrograde traffic to the Golgi (129). Although cellular pIRF3 responses to initial infection have yet to be investigated, it is tempting to speculate that they too may hide the genome from cytoplasmic DNA sensors. The polyomaviruses (PyVs) are dsDNA viruses that are structurally similar to papillomaviruses. PyVs also undergo subcellular retrograde transport after cellular uptake, but unlike HPVs, SV40 and the related BKPyV and JCPyV bypass the Golgi and traffic directly from endosomes to the ER, where redox-dependent chaperones loosen the VP1 capsid to expose membrane-interacting minor capsid proteins VP2/3 (130-132). The post-ER fate of the PyV vDNA is unclear, with data suggesting cell- and virus-type specific differences in transport of vDNA from ER to cytosol/nucleus (133). A recent study found a striking lack of pIRF3 and downstream IFN/ISG responses to BKPyV infection (134), similar to what we observe with papillomaviruses in the present manuscript.

Of note, these smaller viruses do not package tegument or core proteins that may be able to counteract cytoplasmic sensors or their downstream effector proteins. Indeed, these viruses, like papillomaviruses, need to avoid triggering these cytoplasmic sensors. This raises the possibility that multiple viral families may have converged on a trafficking-dependent immunoevasive strategy.

## Materials and Methods

### Tissue Culture

HaCaT cells (82) were grown in high glucose DMEM supplemented with 10% FBS and Ab/Am. Cells were maintained at 37°C with 5% CO_2_ and passaged every 2-3 days. Murine J2 fibroblasts were grown in high glucose DMEM supplemented with 10% NCS, 1% penicillin/streptomycin and 1% L-glutamine. Cells were maintained at 37°C with 5% CO2 and passaged every 3-4 days. To create a feeder layer to support the growth of primary human foreskin keratinocytes, J2 fibroblasts were irradiated every 3-4 days. Cells were removed from continuing culture flasks, resuspended in media in a 15 mL conical, and irradiated with 6000 rads using a Gammacell 40 cesium-137 source as described (135). Irradiated cells were plated at 1 million cells per 10 cm plate to later be used for support of primary keratinocytes. Primary human foreskin keratinocytes (HFKs) were derived from foreskin samples as previously described (136). For continuing culture only, primary HFKs were plated on top of the irradiated J2 fibroblast layer. After trypsinizing or thawing, primary HFKs were grown in high glucose DMEM and F12 media supplemented with 10% FBS, 0.4 μg/mL hydrocortisone, 5 μg/mL insulin, 8.4 ng/mL cholera toxin, 24 μg/mL adenine, and 1% L-glutamine. One day after plating, the media was changed to the same as above but with 5% FBS, 10 ng/mL EGF, and 1% penicillin/streptomycin. The media was supplemented with 10 μM Y-27632 dihydrochloride (Chemdea CD0141) to prevent cellular differentiation and senescence (137). Cells were maintained at 37°C with 10% CO_2_ and passaged every 3-4 days. Y-27632 dihydrochloride was omitted when HFKs were plated for use in experiments.

### Nucleic Acid Transfections

HaCaTs were plated at 60,000 cells per well or HFKs were plated at 90,000 cells per well in a 24-well plate in 500 μL complete media. Cells were transfected with 250 or 500 ng endotoxin-free pGL3 using Lipofectamine 2000 (ThermoFisher 11668) in OptiMEM (Life Technologies). Complexes were made in the following manner: 50 μL OptiMEM was combined with pGL3 (either 250 or 500 ng) or water, and a separate 50 μL of OptiMEM was combined with 2 μL Lipofectamine 2000. Each solution was vortexed, combined, vortexed, and incubated for 15 min at room temperature, prior to dropwise addition of 100 μL of complex to the cells. At various timepoints post-transfection, keratinocytes were washed once with PBS and lysed in 1x RIPA lysis buffer (50 mM Tris-HCl pH 8.0, 150 mM NaCl, 1% NP40, 0.5% sodium deoxycholate, 0.1% SDS), supplemented with 1x reducing SDS-PAGE loading buffer, 1x protease inhibitor cocktail (Sigma P1860), 1mM PMSF and 1x PhosSTOP phosphatase inhibitor cocktail (Roche 04906845001). Samples were then boiled for 5 minutes at 95°C and stored at -80°C until gel electrophoresis.

### HPV16 Production

The R302/5A mutant was generated by site-directed mutagenesis of pXULL-based constructs using the QuikChange XL-II kit and verified by Sanger sequencing (138). Luciferase expressing wt and R302/5A mutant HPV16 virions were generated as previously described (138). Briefly, 293TT cells were CaCl2 co-transfected with the appropriate pXULL based plasmids and the luciferase reporter plasmid pGL3; virus was then purified by CsCl gradient. Encapsidated pGL3 content (viral genome equivalent, vge) was determined by SYBR green qPCR against a standard curve dilution series using primers qLuc2-A (ACGATTTTGTGCCAGAGTCC) and qLuc2-B (TATGAGGCAGAGCGACACC), specific for the luciferase gene in the pGL3 plasmid. DNA concentration of the purified virus was determined by measurement of OD260 on a nanodrop spectrophotometer. From this, the capsid content was calculated and the capsid:pGL3 ratios were determined. HPVs do not utilize specific vDNA packaging signals, and the 293TT method used to generate pGL3-containing HPV16 virions results in the promiscuous packaging of chromatinized cellular DNA, in addition to chromatinized pGL3 reporter plasmid (87, 139). The calculated packaging ratio of 11.16 was therefore used to normalize luciferase data in experiments comparing pGL3 delivery via cationic lipid transfection to wildtype HPV16 infection (Figure 1 and 2).

### HPV16 Infections

HaCaTs or HFKs were plated at 60,000 or 90,000 cells per well, respectively, in a 24-well plate in 500 μL complete media. Cells were infected the following day with wildtype or R302/5A mutant HPV16 virions at 250 ng or 500 ng vDNA equivalents per well for pIRF3 blotting experiments. Infections were performed at 2e8 vge/well for luciferase experiments. For infection experiments with cationic lipids, HPV16 virions were diluted into 50 μL OptiMEM, and a separate 50 μL of OptiMEM was combined with 3 μL Lipofectamine 2000. Solutions were vortexed, combined, vortexed again and incubated for 15 min at RT prior to addition of the 100 μL onto subconfluent HaCaT or HFK cells plated in 500 μL of the appropriate complete media. Nanopure H_2_O was substituted for Lipofectamine 2000 as a control in these experiments. For biochemical inhibition experiments the endosomal acidification and H^+^-ATPase inhibitor Bafilomycin A (BafA, Millipore 196000) was used at 100 nM, furin inhibitor decanoyl-RVKR-cmk (dRVKR, Millipore 344930) was used at 25 μM, and γ-secretase inhibitor XXI (XXI, Millipore 565790) was used at 200 nM, as previously described (27). Luciferase assays were performed 24 hr post-infection.

### Luciferase Assay

HPV infected or pGL3 transfected cells were washed once with PBS and lysed in 100 μL reporter lysis buffer (Promega E3971). Luciferase activity was measured on a DTX-800 multimode plate reader (Beckman Coulter) using Luciferase Assay Reagent (Promega E4550). A fraction of each lysate was blotted for GAPDH to normalize luciferase activity.

### SDS-PAGE & Western Blotting

Samples were resolved by SDS-PAGE and transferred onto a 0.45 μm nitrocellulose membrane. Rabbit monoclonal anti-GAPDH (Cell Signaling 2118, 1:5000), mouse monoclonal anti-IRF3 (Abcam 50772, 1:100), and rabbit monoclonal anti-STING (Cell Signaling 13647, 1:1000) blots were blocked in 5% non-fat powdered milk dissolved in Tris-buffered saline containing 0.1% Tween (TBST). Rabbit monoclonal anti-phospho-IRF3 (Cell Signaling 4947, 1:1000) and rabbit monoclonal anti-phospho-STING (Cell Signaling 19781, 1:1000) blots were blocked in 100% Odyssey blocking buffer (Licor 927–40000). Goat anti-rabbit DyLight 680 (Pierce 35568), goat anti-mouse DyLight 680 (Pierce 35518), goat anti-rabbit DyLight 800 (Pierce 535571) and goat anti-mouse DyLight 800 (Pierce 35521) were used as secondary antibodies at 1:10,000 in either 50% Odyssey blocking buffer/TBST or 5% milk/TBST. Blots were imaged with the Licor Odyssey Infrared Imaging System. Band intensities were quantified by densitometry using ImageJ v1.52a. Briefly, high resolution images were inverted and bands (both pIRF3 and GAPDH for each blot) were carefully boxed to measure the area and mean intensity of each band. An average background intensity value for each blot was used with the appropriate boxed area to subtract background for each specific band. From these values, pIRF3/GAPDH ratios were calculated for each timepoint/condition. Vehicle/VSB control values for each data series were set to “1.00” and relative pIRF3 values were plotted with Prism software.

### RNA-seq Analysis

RNA was isolated from transfected and infected HFKs using Qiagen’s RNeasy Mini Kit (Qiagen 74104). RNA was eluted in 60 μL nuclease-free water, further purified with a TURBO DNA-free Kit (Life Technologies AM1907), then shipped on dry ice to Novogene Corporation (Sacramento, CA). High-quality eukaryotic mRNA libraries were prepared and sequenced on an Illumina platform, yielding >20 million paired-end reads (2×150 bp length) per sample. High-throughput sequencing data were initially processed using the CyVerse Discovery Environment, where FastQC was used for quality screening, HISAT2 (140) for mapping sequences to the human reference genome GRCh38, and featureCounts (141) for generating gene-level read counts (Data Set S1). Differential expression analysis of read counts, including library-size correction and statistical testing (accounting for batch effects with the DESeq model, as specified in Data Set S2), was performed using the DESeq2 Bioconductor package (142) implemented in R (143) via RStudio’s integrated development environment (144). Lists of differentially expressed genes (Data Set S3), with an adjusted *P*-value < 0.05, were analyzed for functional enrichment (Data Set S4) using g:Profiler (145). Heatmaps and volcano plots where constructed in R using the pheatmap (146) and EnhancedVolcano (147) packages. Microsoft Office Suite and Adobe Illustrator software were used to create and compile additional figures.

### Transmission Electron Microscopy

HPV16 samples were prepared with cationic lipids as described above. Each sample was applied to an ultra-thin carbon film over Lacey Carbon Support Film on 400-mesh copper grids (Ted Pella, Inc.) that were glow discharged for 1 min. Excess solution was blotted with Whatman #1 filter paper and the grid was rapidly stained with 2% uranyl acetate. The uranyl acetate was immediately blotted with Whatman #1 filter paper and rapidly stained with 2% methylamine tungstate. The second stain was immediately blotted away with Whatman #1 filter paper and allowed to air dry. Data were collected on a JEOL 3200FS microscope operated at 300 kV. Images were acquired at a magnification of 87,000x and 160,000x, and at 1.0 μm underfocus using a Gatan UltraScan 4000 charge-coupled device (CCD) camera.

### Statistics

Statistical analyses were performed using Prism 6 (GraphPad Software). Significant differences were determined by one-way ANOVA followed by Dunnett’s or Tukey’s multiple comparisons test or two-way ANOVA followed by Sidak’s multiple comparisons test. An alpha of 0.05 was used to determine the statistical significance of the null-hypothesis.

## Acknowledgments

We thank Anne Cress for the HaCaT cells and Jana Jandova for isolation and initial culturing of the primary HFKs. We thank Tony Day of the UA ARL Imaging Core-Life Sciences North for assistance with TEM. We thank Matthew Bronnimann for generating and purifying wildtype HPV16 virions. We are thankful for the bioinformatics tools in the CyVerse Discovery Environment (www.cyverse.org), a computational infrastructure supported by the National Science Foundation under award numbers DBI-0735191, DBI-1265383, and DBI-1743442.

## Supplementary Material

***Figure S1. HPV Virion Infection Does Not Activate Cellular Responses to DNA after 24hr*** Aggregate gene responses following pGL3 DNA introduced via liposome transfection (red, *n* = 2 biological replicates) or HPV virion infection (black, *n* = 3 for Vehicle, 4hr, and 8hr). Preliminary data for 24hr post-infection (blue background, *n* = 1) indicate cellular responses to DNA remain inactive throughout the initial infection. Metagene log_2_ fold change values were computed by aggregating RNA-seq data of all genes within each distinct cellular response cluster (see Figure 3C, D). Error bars represent 95% confidence intervals within each gene cluster.

### Data Set S1

Gene-level read counts, based on the human reference genome GRCh38, for all RNA-seq samples (*n* = 13). Tab-delimited, TXT file, 30.8 MB: “evasion_hg_counts.txt”, available from https://github.com/KVDlab/Uhlorn-2020-evasion

### Data Set S2

RNA-seq sample metadata including batch effects and experimental design groups. Comma-separated, CSV file, 1 KB: “evasion_hg_groups.csv”, available at https://github.com/KVDlab/Uhlorn-2020-evasion

### Data Set S3

Differentially expressed genes, with gene symbol and description annotations, as well as the full differential gene expression results in additional sheets. XLSX file, 9.7 MB: “evasion_hg_DEGs.xlsx”, available at https://github.com/KVDlab/Uhlorn-2020-evasion

### Data Set S4

Functional enrichment analysis of differentially-expressed genes. XLSX file, 0.1 MB: “evasion_hg_functional.xlsx”, available at https://github.com/KVDlab/Uhlorn-2020-evasion

### Data Set S5

Clusters of cellular genes up-regulated in response to DNA. XLSX file, 0.2 MB: “evasion_hg_clusters.xlsx”, available at https://github.com/KVDlab/Uhlorn-2020-evasion

