## Supplementary figures and images for "Vesicular Trafficking Permits Evasion of cGAS/STING Surveillance During Initial Human Papillomavirus Infection"

### Figure S1

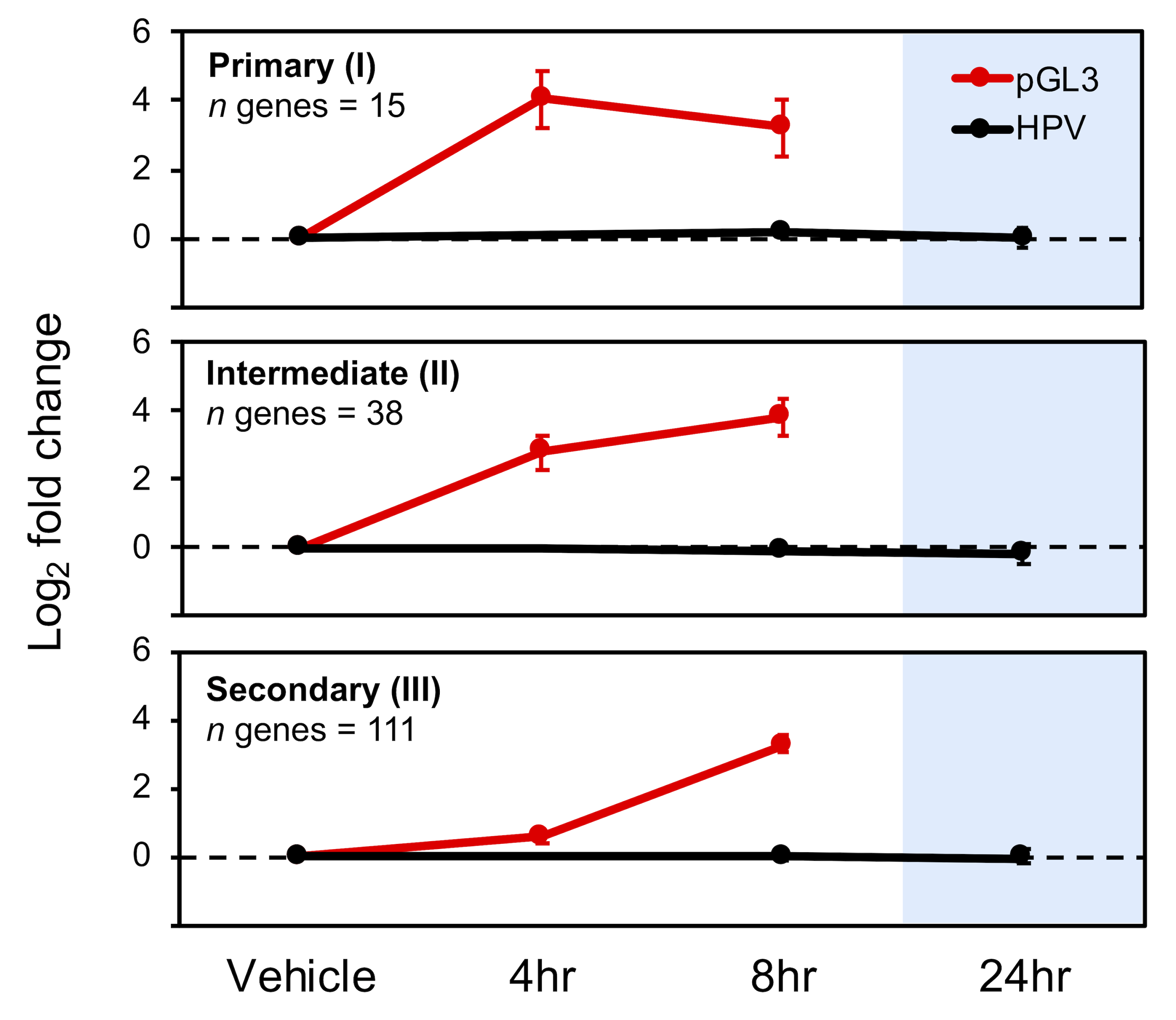
